# uORF-targeting steric block antisense oligonucleotides do not reproducibly activate RNASEH1 expression

**DOI:** 10.1101/2024.06.14.598998

**Authors:** Nina Ahlskog, Nenad Svrzikapa, Rushdie Abuhamdah, Mahnseok Kye, Yahya Jad, Ning Feng, Britt Hanson, Matthew J.A. Wood, Thomas C. Roberts

## Abstract

Upstream open reading frames (uORFs) are *cis*-regulatory motifs that are predicted to occur in the 5ʹ untranslated region (UTR) of the majority of human protein-coding transcripts. uORFs are typically associated with repression of the downstream primary open reading frame (pORF) at either the level of translation, or by promoting mRNA turnover via the nonsense-mediated decay pathway. Interference with uORF activity provides a potential mechanism for targeted upregulation of the expression of specific transcripts. It was recently reported that steric block antisense oligonucleotides (ASOs) can bind to and mask uORF start codons in order to inhibit translation initiation, and thereby disrupt uORF-mediated gene regulation. Given the relative maturity of the oligonucleotide field, such a uORF blocking mechanism might have widespread therapeutic utility. Here, we re-synthesised three of the most potent ASOs targeting the *RNASEH1* uORF described in the study by Liang *et al*. and investigated their potential for RNASEH1 protein upregulation. No upregulation (of endogenous or reporter protein expression) was observed with any of the oligonucleotides tested at doses ranging from 25 nM to 300 nM. Conversely, we observed downregulation of expression in some instances, consistent with well-established mechanisms of blocking ribosome procession. Experiments were performed using multiple transfection protocol setups, with care taken to replicate the conditions of the original study. Transfection efficiency was confirmed using a *MALAT1*-targeting gapmer ASO as a positive control. We conclude that previously-described *RNASEH1* uORF-targeting steric block ASOs are incapable of upregulating pORF protein expression in our hands.

## Introduction

Steric block oligonucleotides are short single-stranded nucleic acid polymers that bind to target nucleic acid molecules via Watson-Crick base pairing in order to interfere with some binding partner interaction. Multiple steric block antisense oligonucleotides (ASOs) have now received regulatory approval for the treatment of Duchenne muscular dystrophy (eteplirsen, viltolarsen, golodirsen, and casimersen) and spinal muscular atrophy (nusinersen) with further approvals likely for these indications, and others.^1^ Decades of pre-clinical and clinical development have established patterns of ASO chemical modification and routes-of-delivery that provide blueprints for the generation of new therapeutics. The promise of such molecular medicines is that by careful sequence design, successful ASO platform chemistries can be directed to different gene targets. This is exemplified by milasen, a steric block ASO (based on the nusinersen template) designed to treat a single patient with neuronal ceroid lipofuscinosis 7 (CLN7).^2^ Steric block ASOs have primarily been used for splice correction (i.e. to induce exon skipping or exon inclusion so as to correct the translation reading frame in otherwise out-of-frame transcripts). However, steric block oligonucleotides have similarly been utilised for multiple other purposes, including splice corruption to disrupt the translation reading frame,^3^ generation of ectopic proteins with novel functions,^4^ targeting polyadenylation signals for modulating differential poly(A) tailing,^5^ disrupting exon-junction complex formation to relieve nonsense-mediated decay,^6^ removal of ‘poison’ exons containing premature termination codons (also known as targeted augmentation of nuclear gene expression, TANGO),^6–8^ inhibition of translation initiation for gene silencing,^9,10^ and the targeting of upstream open reading frames (uORFs).^11^

uORFs consist of a start codon located within the 5ʹ untranslated region (5ʹ UTR) of messenger RNA (mRNA), followed by an in-frame stop codon. uORFs are common in mammalian transcripts, and are typically associated with translational repression of the downstream primary open reading frame (pORF).^12^ In 2016, a team from Ionis Pharmaceuticals (Liang *et al*.,) reported activation of four genes (human: *RNASEH1*, *SFXN3*, and murine: *Mrpl11*, *Lrpprc*) using a variety of uORF-targeting steric block ASOs.^11^ There is currently a paucity of technologies capable of targeted activation of specific genes. As such, this study was important, as it suggested that relief of uORF-mediated repression could be utilised as a widely-applicable means of therapeutic gene upregulation. To this end, we were motivated to explore the potential of uORF-targeting ASOs. Here we report extensive efforts to reproduce the findings reported by Liang *et al*.^11^ We conclude that previously-described steric block ASOs do not reproducibly activate RNASEH1 expression via uORF start codon masking.

## Results

### uORF-targeting ASOs

Firstly, we selected three oligonucleotides that exhibited the highest degree of RNASEH1 protein upregulation as reported by Liang *et al*.^11^ These molecules consisted of (i) a 16mer phosphodiester 2ʹ-*O*-methyl RNA (PO-2OMe), (ii) a 16mer phosphorothioate 2ʹ-*O*-methoxyethyl RNA (PS-MOE), and (iii) an 18mer phosphorothioate 2ʹ-*O*-methyl RNA (PS-2OMe) (**Figure 1A**). These ASOs are referred to as 761909, 759304, and 783679 in the Liang *et al*., study, respectively.^11^ These oligonucleotides consisted of the same sequence (with the exception of the 18mer which included an additional 2 nucleotides at the 3ʹ terminus). Chemistry controls were synthesised in parallel (**Figure 1A**), in which the constituent nucleotide sequence was scrambled while the patterns of each chemistry were maintained (**Figure 1B**). A gapmer ASO targeting the long non-coding RNA (lncRNA) *MALAT1*^13,14^ was synthesised as a control for transfection (**Figure 1A**). The target *RNASEH1* transcript (NM_002936.6) contains one uORF which overlaps with the pORF and encodes a nine amino acid peptide (**Figure 1C**). The 5ʹ ends of the uORF-targeting ASO sequences are complementary to the start codon of the *RNASEH1* uORF (**Figure 1C**). Publicly available riboseq data^15^ showed a footprint of ribosome occupancy at the *RNASEH1* uORF, and a pronounced initiating ribosome peak at the corresponding upstream ATG (**Figure 1C**). These data suggest that this sequence is a *bona fide* uORF. The relative lack of ribosome initiation at the pORF ATG suggests that the *RNASEH1* is likely subject to uORF-mediated translational repression.

**Figure 1.**
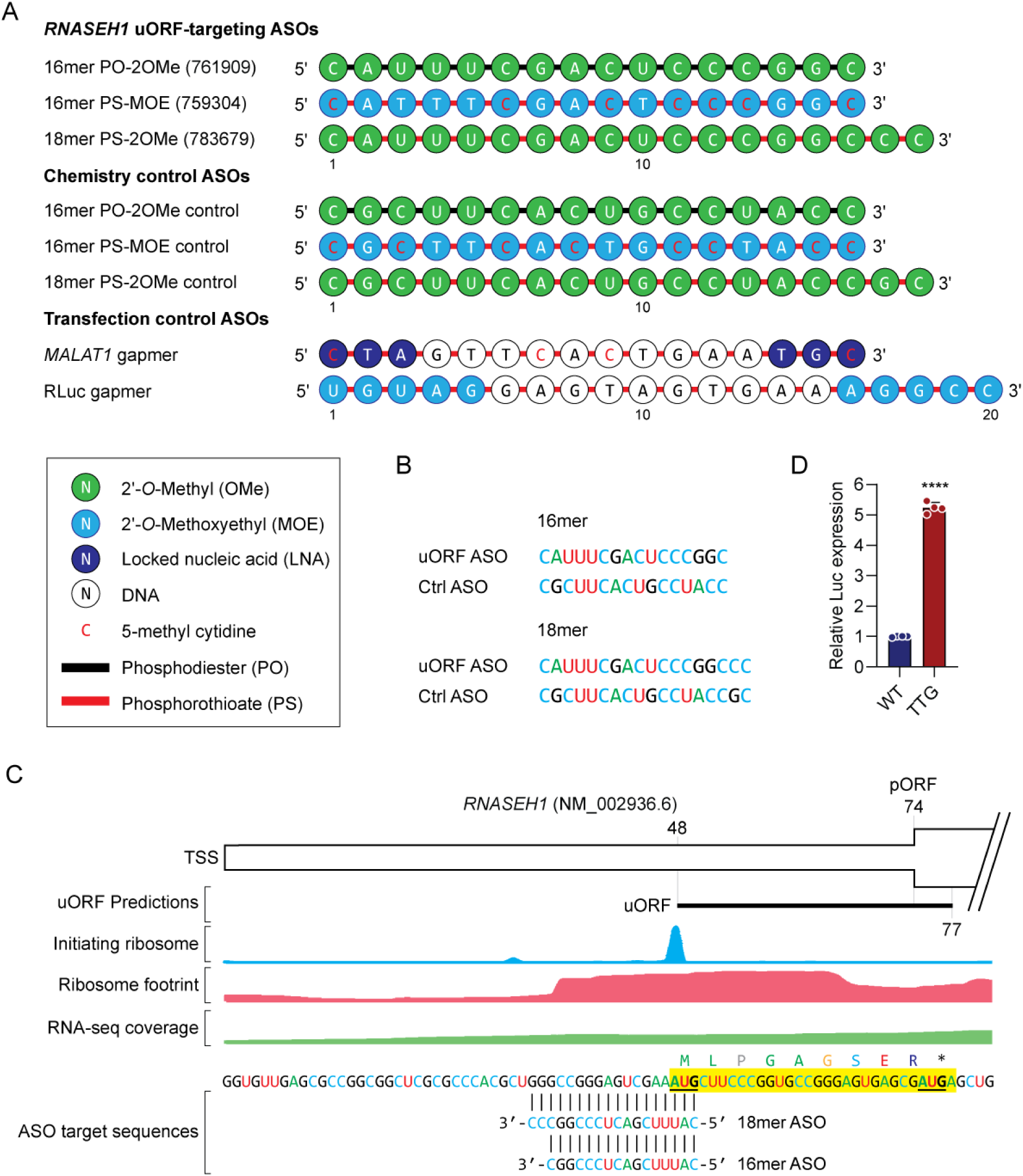
*RNASEH1* uORF-targeting steric block antisense oligonucleotides. (**A**) Schematic of ASO sequences and chemistries used in this study. (**B**) Sequences of on-target *RNASEH1* uORF-targeting ASOs and their scrambled control (Ctrl) sequences for the 16mer and 18mer variants. (**C**) Schematic of the *RNASEH1* transcript showing the position, sequence, and translation of the uORF and binding locations for the uORF-targeting ASOs. Aggregated riboseq and RNA-seq data are overlaid providing evidence of uORF translation. The uORF is highlighted in yellow and the start codons are highlighted in bold and underlined. TSS, transcription start site, pORF, primary open reading frame. (**D**) HEK293T cells were transfected with *RNASEH1* 5ʹ UTR dual luciferase reporter constructs as indicated and luciferase activity assayed after 24 hours. A mutant construct (TTG) in which the predicted uORF was disrupted by mutation of the uORF start codon was analysed in parallel. Values are mean+SEM (*n*=4), and were scaled such that the mean of the WT control group was returned to a value of 1. Statistical significance was determined by unpaired Student’s *t*-test, \*\*\*\**P*<0.0001.

Dual luciferase reporter (DLR) constructs were generated in which the *RNASEH1* 5ʹ UTR was cloned upstream of a Renilla luciferase transgene. A mutant construct in which the *RNASEH1* uORF start codon was ablated (by mutating its ATG start codon to TTG) was generated in parallel. These constructs were transfected into HEK293T cells and relative luciferase activity determined 24 hours later. A pronounced increase (>5-fold, *P*<0.0001) in Renilla luciferase signal was observed for the mutant ‘TTG’ construct, thereby validating that the *RNASEH1* uORF does indeed repress its downstream pORF (**Figure 1D**), consistent with findings reported by Liang *et al*.^11^

### *RNASEH1* uORF-targeting ASOs do not activate endogenous protein expression

We next sought to determine whether uORF-targeting ASOs could influence endogenous RNASEH1 protein expression. HeLa cells were transfected with 100 nM of each of the uORF-targeting ASOs (or chemistry controls) and protein harvested over a range of time points. RNASEH1 protein upregulation effects were previously reported at 5, 10, 12, 16, and 24 hours post transfection by Liang *et al*.^11^ We expanded this range by collecting protein lysates at 4, 8, 12, 24, 48, and 72 hours post transfection. Protein lysates were analysed using the Jess capillary western system using a commercially-available anti-RNASEH1 antibody. (Notably, the anti-RNASEH1 antibody used in the Liang *et al*., study was developed in-house and so was not available to us). Vinculin (VCL) was used as a loading control, which was highly consistent between samples. No increase in RNASEH1 protein expression as observed for any of the ASO treatments relative to either untreated cells or matched chemistry controls at any time point for *n*=4 completely independent experiments (**Figure 2,3**). RNASEH1 western blot signal was observed at the expected size (32 kDa, 268 amino acids), and the specificity of the anti-RNASEH1 antibody was confirmed by siRNA knockdown (**Figure S1**). Very similar data were obtained using conventional SDS-PAGE western blots for the 24 and 48 hours post transfection lysates, whereby equal protein loading was assessed by both VCL immunoblotting and Fast Green membrane staining for total protein loading (**Figure S2**). Parallel transfection of an ASO gapmer targeting the ubiquitously-expressed lncRNA *MALAT1* resulted in target knockdown that was statistically significant (*P*<0.05) at 24 hours post transfection and which reached a peak of >75% target knockdown relative to untreated control cultures at 72 hours post transfection (*P*<0.0001), thereby excluding poor transfection as an explanation for a lack of RNASEH1 activation effect (**Figure S3**).

**Figure 2.**
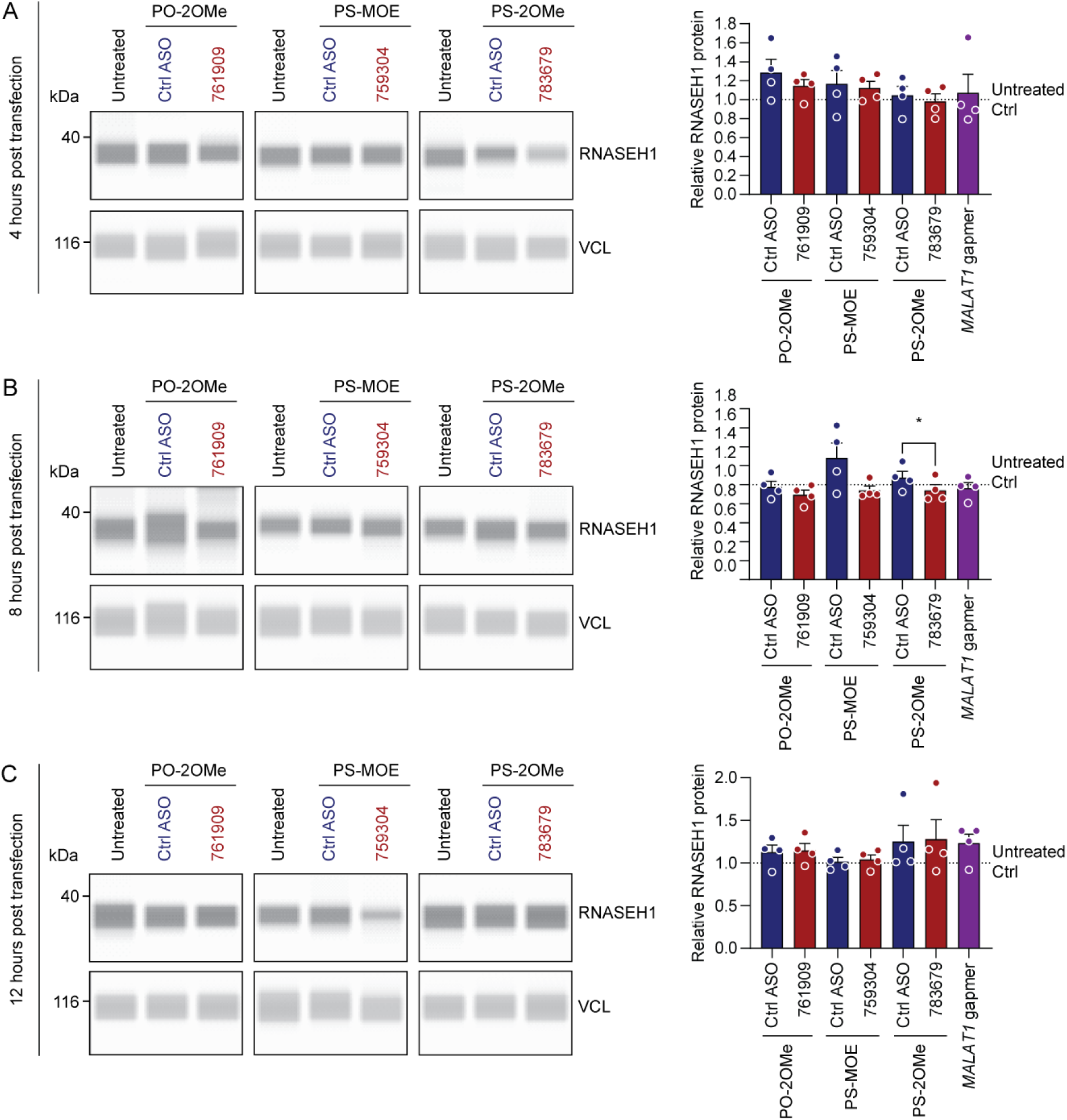
uORF-targeting steric block ASOs do not increase RNASEH1 protein expression at 4, 8, and 12 hours post transfection. HeLa cells were transfected with 100 nM ASOs or matched chemistry controls and cells harvested at (**A**) 4 hours, (**B**) 8 hours, or (**C**) 12 hours post transfection. RNASEH1 protein was quantified by Jess capillary western blot. Vinculin (VCL) was used as a loading control. Representative blots are shown together with histograms of protein quantification. The values of untreated control samples are indicated by dotted lines (scaled to a value of 1). A gapmer targeting *MALAT1* was included as a positive control for transfection, which is not expected to influence RNASEH1 expression. Values are mean+SEM. Statistical significance was assessed by paired Student’s *t*-test between each treatment and its respective control ASO, \**P*<0.05, *n*=4 completely independent experiments.

**Figure 3.**
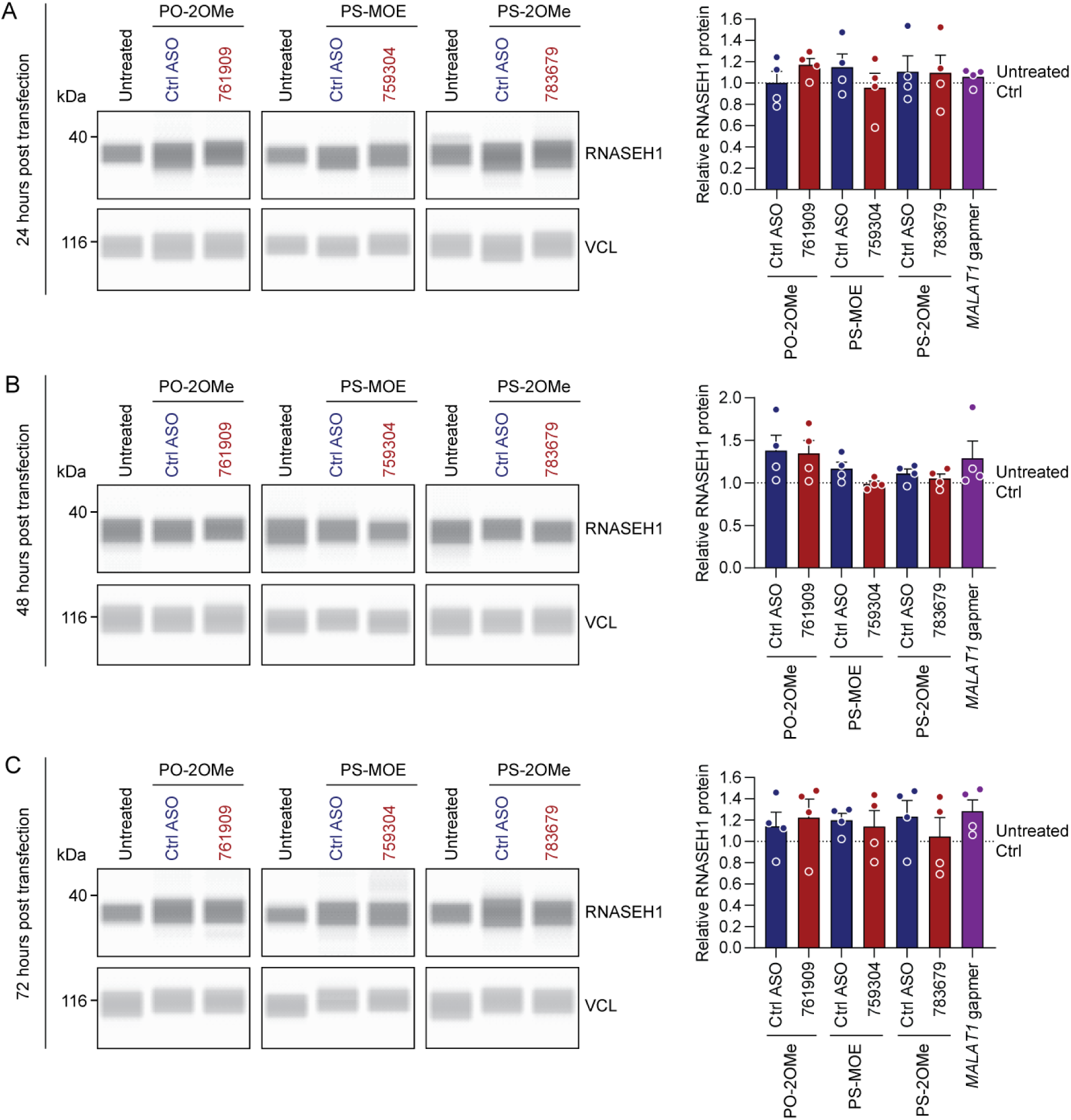
uORF-targeting steric block ASOs do not increase RNASEH1 protein expression at 24, 48, and 72 hours post transfection. HeLa cells were transfected with 100 nM ASOs or matched chemistry controls and cells harvested at (**A**) 24 hours, (**B**) 48 hours, or (**C**) 72 hours post transfection. RNASEH1 protein was quantified by Jess capillary western blot. Vinculin (VCL) was used as a loading control. Representative blots are shown together with histograms of protein quantification. The value of untreated control samples is indicated by the dotted line (scaled to a value of 1). A gapmer targeting *MALAT1* was included as a positive control for transfection, which is not expected to influence RNASEH1 expression. Values are mean+SEM. Statistical significance was assessed by paired Student’s *t*-test between each treatment and its respective control ASO, (no significant changes detected), *n*=4 completely independent experiments.

### *RNASEH1* uORF-targeting ASOs do not activate endogenous protein expression at doses ranging from 25 to 300 nM

Protein upregulation effects were reported at concentrations ranging from 20 nM to 100 nM by Liang *et al*.^11^ Interestingly, the same study reported parabolic dose responses in some cases, or no dose response in others.^11^ As such, we reasoned that potential protein upregulation responses might be observed at some concentrations but not others. We therefore conducted a dose response experiment in HeLa cells with all three uORF-targeting ASO chemistries (and their matched chemistry controls) at 25, 50, 100, 200, and 300 nM. Protein lysates were harvested 48 hours post transfection. No upregulation was observed for any oligonucleotide chemistry at any concentration tested (**Figure 4**). By contrast, treatment with PS-MOE ASOs resulted in RNASEH1 downregulation, which reached statistical significance at the 25, 50, 100, and 200 nM concentrations (**Figure 4B**). Treatment with the PS-2OMe ASO also reduced RNASEH1 protein expression at the 300 nM concentration, although this effect did not reach statistical significance at the *P*<0.05 level. Results were obtained for *n*=4 completely independent experiments (and *n*=8 independent experiments for the 100 nM concentration). The validity of the transfection protocol was confirmed for these experiments using the *MALAT1*-targeting ASO gapmer, which exhibited ∼75% knockdown relative to untreated controls, and the PS-2OMe or PS-MOE control ASOs (**Figure 4D**, *P*<0.001).

**Figure 4.**
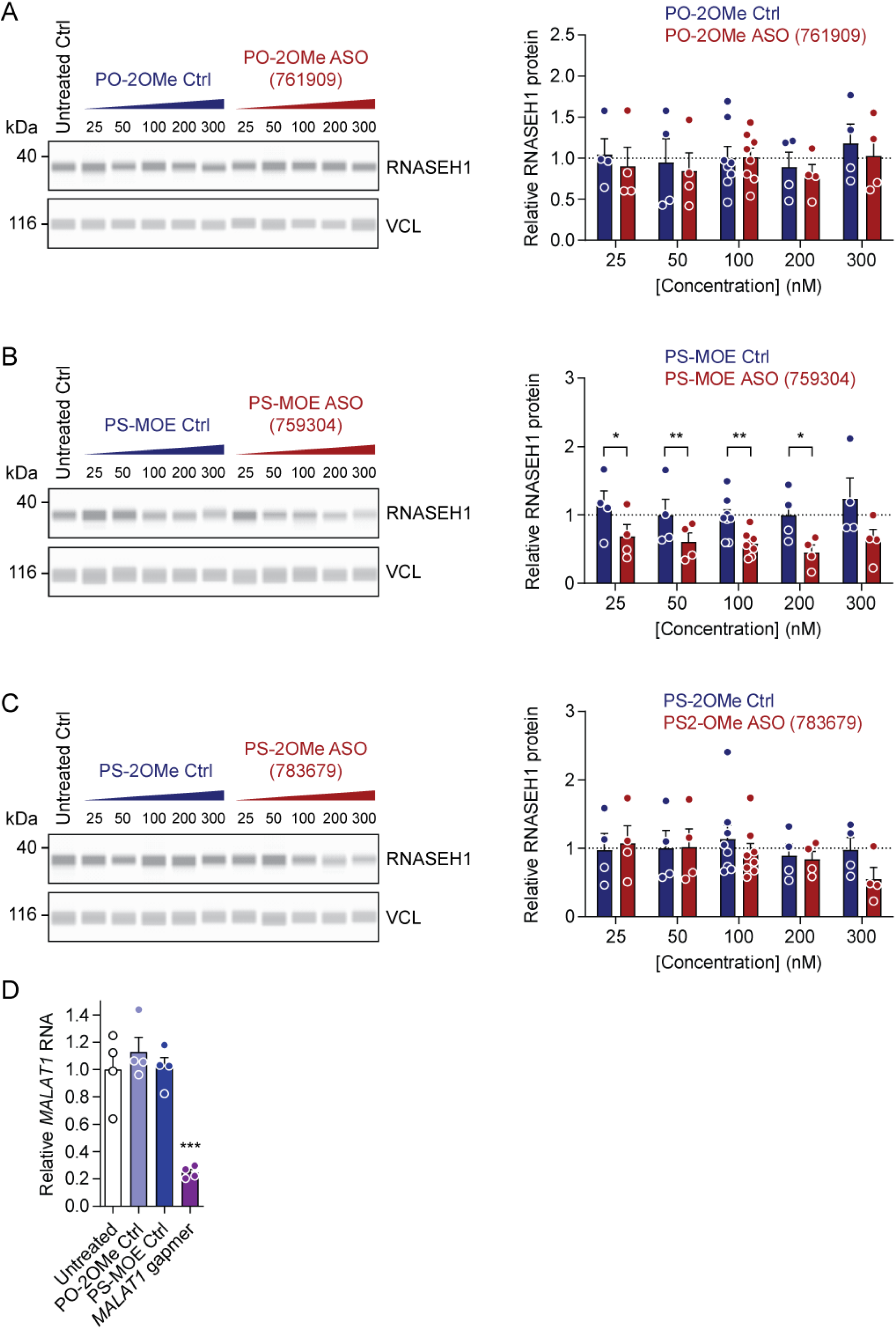
uORF-targeting steric block ASOs do not increase RNASEH1 protein expression regardless of dose. HeLa cells were transfected with ASOs at concentrations as indicated and protein harvested after 48 hours for (**A**) PO-2OMe, (**B**) PS-MOE, and (**C**) PS-2OMe nucleic acid chemistries. RNASEH1 protein was quantified by Jess capillary western blot. Vinculin (VCL) was used as a loading control. Representative blots are shown together with histograms of protein quantification. The value of untreated control samples is indicated by the dotted line (scaled to a value of 1). (**D**) Cells were transfected with a gapmer targeting *MALAT1* (100 nM) in parallel as a positive control for transfection. *MALAT1* transcript levels were determined by RT-qPCR and normalized to *RPL10* expression. Values are mean+SEM. Statistical significance for protein data were assessed by paired Student’s *t*-test within each oligonucleotide dose. RT-qPCR data were analysed by one-way ANOVA and Tukey *post hoc* test. \**P*<0.05, \*\**P*<0.01, \*\*\**P*<0.001, *n*=4 or 8 independent experiments as indicated.

### uORF-targeting ASOs do not activate *RNASEH1* 5ʹ UTR luciferase reporters

Liang *et al*. previously reported that uORF-targeting ASOs can activate luciferase reporter constructs.^11^ We therefore transfected HeLa cells with *RNASEH1* 5ʹ UTR-DLR reporter plasmids, followed by a second transfection with uORF-targeted ASOs and luciferase activity determined 24 hours later (**Figure 5A**). ASOs were transfected at final concentrations of 100 nM or 50 nM together with matched chemistry controls. An ASO gapmer targeting RLuc was utilised as a positive control for transfection. Transfection of the RNASEH1-TTG mutant plasmid (with the uORF start codon disrupted) was included as an additional control intended to demonstrate the theoretical maximum RNASEH1 upregulation effect. No significant changes were observed for any uORF-targeting ASO relative to the untreated control, or to any of the matched chemistry controls at either dose (**Figure 5B**). The positive control RLuc gapmer significantly (*P*<0.05) reduced target expression by ∼66%, indicative of a robust dual transfection protocol (**Figure 5B**, *n*=5 independent experiments). Very similar results were observed when the same experiment was performed in HEK293T cells (**Figure 5C**, *n*=3 independent experiments). These data suggest that previously-described uORF-targeting steric block ASOs do not activate *RNASEH1* 5ʹ UTR report constructs.

**Figure 5.**
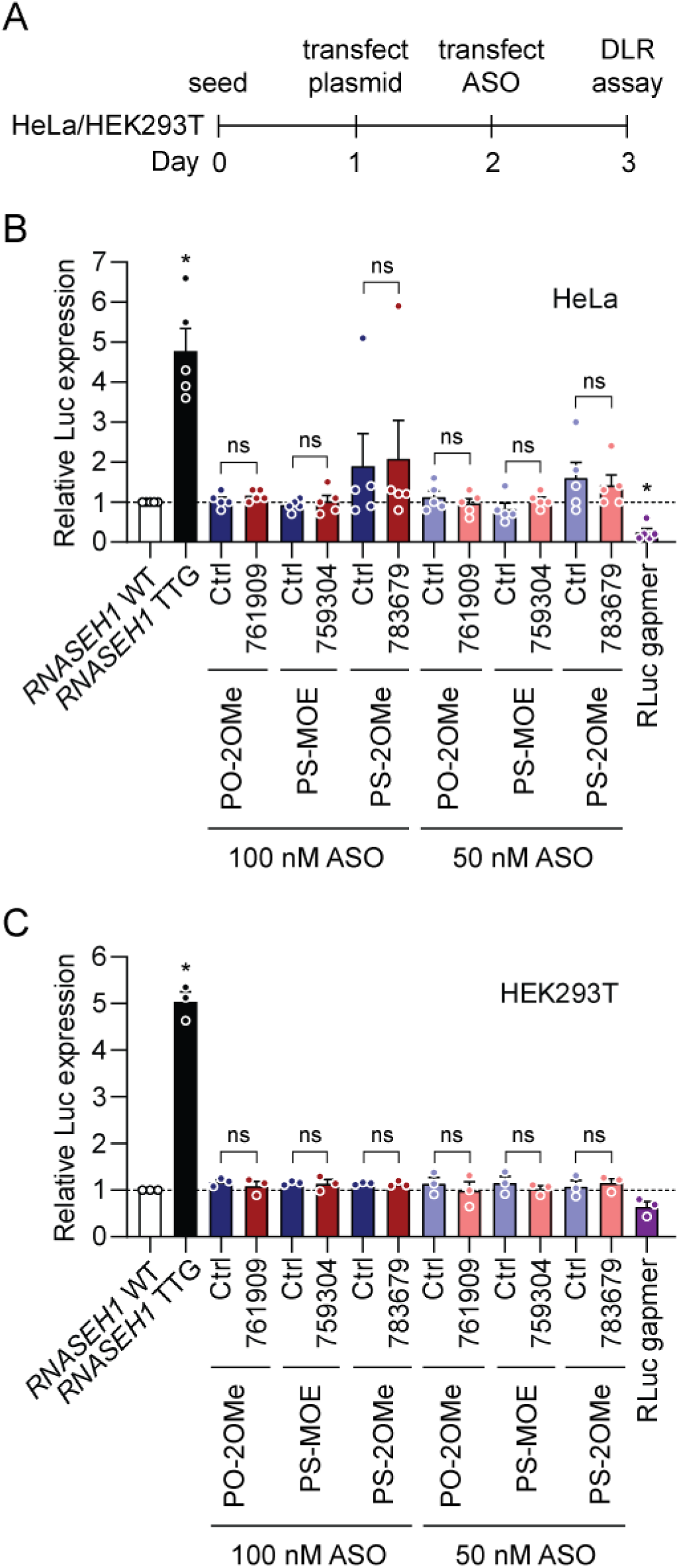
uORF-targeting steric block ASOs do not activate a *RNASEH1* 5ʹ UTR driven luciferase reporter construct. (**A**) Schematic of experimental design for sequential plasmid and ASO transfection. Cells were first transfected with plasmids encoding *RNASEH1* 5ʹ UTR dual luciferase reporter (DLR) constructs. After 24 hours, cells were transfected with ASOs as indicated. Cells were subsequently harvested after a further 24 hours. Renilla luciferase activity was determined for *RNASEH1* 5ʹ UTR constructs and signal normalised to firefly luciferase (encoded from a cistronically-independent transgene cassette) for (**B**) HeLa cells (*n*=5 independent experiments), and (**C**) HEK293T cells (*n*=3 independent experiments). A gapmer targeting RLuc (100 nM) was used as a positive control for transfection. A TTG mutant in which the *RNASEH1* uORF start codon is disrupted indicated the theoretical maximum of reporter upregulation. Values are mean+SEM. Statistical significance was assessed by one-way ANOVA and Tukey *post hoc* test, ns, not significant, \**P*<0.05. Statistical comparisons are to the *RNASEH1* WT group unless otherwise indicated.

### Confirmation of ASO integrity

We performed a series of analyses to exclude the possibility that the integrity of ASOs used in the present study had somehow been compromised. The integrity of each ASO was assessed by MALDI-TOF-MS, whereby prominent, single m/z peaks were observed for all oligonucleotides (with double-ionized peaks also available in some instances) **(Figure S4**). The observed m/z peak values for each ASO were within less than 0.2% of the expected mass. Similarly, ASOs were analysed by LC-MS, whereby prominent single peaks were observed for each oligonucleotide with an observed mass within 0.2% of the expected mass (and all but one ASO was within >0.02%) (**Figure S5**). These data confirm the integrity of the ASOs used in this study.

## Discussion

Using the three most potent uORF-targeting steric block oligonucleotides described by Liang *et al*.^11^ we were unable to observe an increase in RNASEH1 protein relative to untreated cultures or cells treated with matched chemistry controls. This was true irrespective of model system, ASO chemistry, ASO length, ASO dose, or collection time point (**Figure 2,3,4,5**). The RNASEH1 antibody was validated as specific (**Figure S1**), and protein quantification was performed using both the Jess capillary western blot system and standard SDS-PAGE western blots (**Figure 2,3,4,S2**). uORF-targeting ASOs were synthesised twice, using a commercial supplier (IDT). Successful ASO transfection was confirmed by simultaneous treatment with a gapmer targeting *MALAT1* (**Figure S3, 4D**). Integrity of ASOs was confirmed by in-house MALDI-TOF-MS and LC-MS analyses (**Figure S4,S5**). Based on these experiments we conclude that we are unable to reproduce the findings of Liang *et al*. with respect to RNASEH1 upregulation reported previously.^11^

Steric block ASOs (and specifically, phosphorodiamidate morpholino oligonucleotides, PMOs) have been extensively used to target the 5ʹ UTRs (usually, but not exclusively, in the vicinity of the pORF start codon) as a gene silencing technology, especially in the zebrafish *Danio rerio*.^9,10,16–18^ Such translation blocking ASOs have been assessed in clinical trials, as in the case of AVI-4126 for the targeting of the *MYC* oncogene.^19^ In these cases, ASO binding is expected to interfere with assembly of the 80S ribosome, and or, prevent ribosome procession. Whether the 40S subunit can continue scanning beyond the site of ASO binding is unknown. However, were that to be the case, then targeting sequences in the 5ʹ UTR (i.e. uORFs) with ASOs might be expected to inhibit both uORF and pORF translation, with the net effect being translational repression of the pORF. Such a mechanism would be consistent with our experimental observations using the PS-MOE ASO chemistry, which induced RNASEH1 down-regulation instead of the intended activation (**Figure 4B**).

At the time of writing (May 2024) the study by Liang *et al*. in *Nature Biotechnology*^11^ has been cited 173 times. A follow-up study by the same authors was published in 2017 in *Nucleic Acids Research*^20^ and has been cited 95 times, with a total 210 unique citations across both articles. The majority of citing articles were reviews (*N*=105, 50%). Research articles constituted 27% of all citations (*N*=56), which were further analysed to determine the extent to which uORF-targeting ASOs have been adopted by other researchers. Three articles were identified as being particularly relevant. For example, Sasaki *et al*., reported upregulation of CFTR using uORF-targeting ASOs.^21^ Furthermore, ASOs targeting a non-uORF translation repression element also induced CFTR protein upregulation. No ASO sequences were provided in this report, meaning that it is not possible to assess the exact target locations of the functional ASOs at this time. Interestingly, in this study, simultaneous disruption of the uORF start codon and a neighbouring structured region was required to induce activation of a downstream reporter gene.^21^ Similarly, Kidwell *et al*., also reported targeting activation of murine *Ppp1r15a* using steric block ASOs which bind to a pORF-proximal uORF.^22^ In this case, a uORF start codon overlapping ASO was found to induce a modest increase in reporter expression, but an ASO that bound downstream and which did not overlap with the uORF start codon induced more robust upregulation (including for endogenous PPP1R15A protein in mouse kidney).^22^

A uORF targeting strategy was also reported by Tan *et al*., using steric block ASOs targeting the 5ʹ UTR of *NUDT21* in the context of kidney renal clear cell carcinoma (KIRC).^23^ While robust NUDT21 protein upregulation was observed, the functional ASOs were actually non-overlapping with the predicted uORF start codon, whereas overlapping ASOs were non-functional, suggestive of a distinct mechanism.^23^

Importantly, a number of the other citing studies reported upregulation effects by targeting 5ʹ UTR sequences with steric block ASO oligonucleotides in a uORF-independent manner. For example, it has been reported that ASOs targeting 5ʹ UTR structural elements can induce targeted protein upregulation.^20^ Steric block ASOs designed to disrupt the interplay between 5ʹ UTR double-stranded RNA motifs and uORF translation were shown to modulate pORF translation.^24^ Steric block ASOs have also been shown to promote increased expression through mRNA stabilisation. Specifically, ASOs targeting the frataxin (*FXR*) 5ʹ UTR stabilised mRNA turnover, leading to increased transcript and protein levels in a uORF-independent manner. Interestingly, the 5ʹ UTR-targeting ASOs in this study bound in close proximity to the pORF start codon, which might otherwise have been expected to exhibit an inhibitory effect on translation.^25^ Similarly, an ASO complementary to a predicted uORF in the *SMN2* transcript resulted in mRNA stabilisation and protein upregulation which was not associated with uORF activity.^26^ These studies suggest that the 5ʹ UTR constitutes a promising site for potential therapeutic oligonucleotides, and that in some cases, binding to uORF sequences may be incidental. The remaining citing research articles did not utilise ASOs targeting uORFs or 5ʹ UTRs, and so were deemed not relevant. In summary, the number of studies reporting uORF start codon-targeting steric block ASOs activating protein expression is very limited. Notably, we are aware that groups from Roche and Astra Zeneca recently presented preliminary data on the topic of uORF targeting for protein upregulation at the Oligonucleotide Therapeutics society 2023 meeting.

While reproducing the upregulation effects reported for RNASEH1 has been challenging, the observation that this transcript is regulated by its corresponding uORF is robust (**Figure 1C,D**). Notably, alternative methods of uORF interference may still provide means of therapeutic gene upregulation, especially for monogenic haploinsufficiency disorders such as Angelman syndrome, Dravet syndrome, and Rett syndrome. The application of alternate technologies such as RNA structure disruption,^24^ synthetic IRES scaffolds,^27^ SINEUP lncRNAs,^28^ and exon skipping of uORF containing exons^29^ are exciting possibilities for future nucleic acid-based therapeutics to treat such disorders. In conclusion, this study casts doubt on the notion that steric block ASOs targeted to a uORF start codon can induce protein upregulation in the specific case of RNASEH1, and possibly to some extent, in the general case.

## Methods

### Oligonucleotides

ASOs were purchased from Integrated DNA Technologies (IDT, Leuven, Belgium) or synthesised in-house (i.e. for *MALAT1* and RLuc gapmer ASOs). All ASO sequences are listed in **Table S1** and illustrated diagrammatically in **Figure 1A**.

Small interfering RNA (siRNA) pools were purchased from Dharmacon (Horizon Discovery LTD, Cambridge, UK). ON-TARGETplus Human RNASEH1 siRNA (246243) (target sites: 5ʹ-GACAGUAUGUUUACGAUAA-3ʹ, 5ʹ-GAGCACAGGUGGACCGGUU-3ʹ, 5ʹ-ACAAGAAUCGGAGGCGAAA-3ʹ, 5ʹ-GAGCAGGAAUCGGCGUUUA-3ʹ). A non-targeting siRNA pool was used as a negative control (D-001810-10-05, Dharmacon).

### Cell cultures

HeLa and HEK293T cells were grown in culture media composed of DMEM-Glutamax supplemented with 10% foetal bovine serum (both Gibco, Thermo Fisher Scientific, Loughborough, UK) and 1% Antibiotic/Antimycotic solution (Merck Life Science, Gillingham, UK). The cells were maintained in a passively humidified incubator at 37°C, 5% CO_2_. Cell cultures were confirmed free of mycoplasma contamination through monthly testing.

ASOs were transfected using Lipofectamine RNAiMAX (Invitrogen) according to manufacturer’s instructions. Plasmid DNA (100 ng per well) was transfected using Lipofectamine 2000 (Invitrogen) according to manufacturer’s instructions.

### Protein quantification by capillary gel electrophoresis

Cells were plated in ∼4×10^5^ cells/well in 2 ml culture media on 6-well plates and transfected 24 hours later with ASOs as above at indicated concentrations. The treated cells were returned to the incubator for 4, 8, 12, 24, 48, or 72 hours. At the specified time point, the cells were collected in 150 µl of ice-cold RIPA buffer (Thermo Fisher Scientific) with cOmplete EDTA-free protease inhibitor (Roche, Welwyn Garden City, UK) and sonicated 3×5 seconds using a Q500 Sonicator (Qsonica LLC, Newtown, CT, USA) at 25% amplitude. The lysates were cleared by centrifugation at 10,000 *g* for 5 minutes and protein concentration was determined using the DC Protein Assay (Bio-Rad Laboratories, Watford, UK) according to the manufacturer’s instructions. All samples were diluted to a final concentration of 1.5 mg/ml total protein in RIPA buffer prior to loading on the Jess instrument or for SDS-PAGE.

Capillary gel electrophoresis experiments were performed using the Jess Simple Western system with a 12-230 kDa fluorescence separation module (both Bio-Techne, Abingdon, UK) according to manufacturer’s instructions. Primary antibodies anti-RNaseH1 (1:250, #15606 Proteintech, Manchester, UK) and anti-Vinculin (1:25,000, #V9131 Sigma-Aldrich, Gillingham, UK) were used for relative quantification and loading normalization purposes respectively. Secondary antibodies αRb-IR and αMs-NIR (both Bio-Techne, Abingdon, UK) were used as per manufacturer’s instructions (i.e. undiluted). The results were analysed using the Compass for Simple Western software (version 6.3.0) according to manufacturer’s instructions.

### RT-qPCR

Total RNA was isolated from the cells of 6-well plates treated as above using the Maxwell RSC simplyRNA kit and the Maxwell RSC Instrument (both Promega, Southampton, UK) according to manufacturer’s instructions. In short, the cells were washed and collected by trypsinization. The cell pellet was then resuspended in 200 µl of Maxwell Homogenization solution containing 1-thioglycerol and loaded onto prefilled Maxwell RSC cartridges and run in Maxwell RSC 16 with the appropriate program. Extracted RNA was quantified by UV spectrophotometry using a Nanodrop 2000 instrument (Thermo Fisher Scientific). Complementary DNA (cDNA) was reverse-transcribed from 1 µg of total RNA using the High-Capacity cDNA Reverse Transcription Kit (Thermo Fisher Scientific) according to manufacturer’s instructions.

Quantitative polymerase chain reaction (qPCR) was performed on the cDNA (diluted 1 in 5 in nuclease-free water) using the Power SYBR Green PCR Master Mix and the StepOnePlus Real-Time PCR System (both Applied Biosystems, Warrington, UK). *MALAT1* expression was normalised to *RPL10* (60S ribosomal protein L10) and relative quantification determined using the ΔΔCt method.^30^ Primer sequences are listed in **Table S2**.

### Dual luciferase reporter assay

For the luciferase reporter assay, cells were plated ∼1.5×10^4^ cells/well in 80 µl of culture media in white-walled 96-well plates. After 24 hours, the cells were transfected with 100 ng/well of Renilla luciferase reporter gene with either a WT or a TTG-mutated 5ʹ UTR for RNASEH1 using the Lipofectamine 2000 Transfection Reagent (Thermo Fisher Scientific). On the following day (48 h), the cells were transfected with either 50 or 100 nM ASO as described above, or 100 nM RLuc gapmer (positive control for transfection). On day three (72 h), the cells were assayed using the Dual-Glo Luciferase Assay System (Promega) and a Clariostar Plus plate reader (BMG Labtech, Aylesbury, UK).

### Western Blot (SDS-PAGE)

Protein samples (20 mg of total protein per lane) were separated by SDS-PAGE using precast 10% NuPAGE Bis-Tris midi gels (Thermo Fisher Scientific). Protein was electroblotted onto PVDF (polyvinylidene fluoride) membrane (Merck Milipore, Watford, UK). Membranes were stained with Fast Green FCF (Sigma-Aldrich) and imaged for total protein on a ChemiDoc MP Imaging System (Bio-Rad). Membranes were blocked in Intercept PBS Blocking Buffer (Li-Cor Biotechnology, Cambridge, UK) before over-night incubation with anti-RNaseH1 (1:500, Proteintech, Manchester, UK) and anti-Vinculin (1:20,000, Sigma-Aldrich) antibodies in blocking buffer. Following incubation with HRP-linked secondary antibody (1:500 Horse αMs-HRP, #7076S or Gt αRb-HRP, #7074S; both Cell Signaling Technology, Leiden, The Netherlands), the signal was developed using Clarity Western ECL Substrate and visualized on the ChemiDoc MP Imaging System (both Bio-Rad).

### MALDI-TOF-MS

For MALDI-TOF-MS analysis, 1 µl of 100 µM ASO was added to 10 µl of MALDI matrix solution (40 mg/ml 3-hydroxypicolinic acid, 40 mM ammonium citrate in acetonitrile/water [1:1]), and mixed thoroughly. The resulting mixture was spotted (0.5 µl) onto a MALDI target plate, dried, and then analysed using a Shimadzu MALDI-8020 instrument (Shimadzu UK Ltd, Milton Keynes, UK).

### LC-MS

LC-MS analysis was performed on a Waters SQD 2 coupled to a Waters ACQUITY UPLC system using a Waters ACQUITY Premier BEH C18 1.7 µm column (2.1×50 mm) (all Waters, Wilmslow, UK). ASO samples were adjusted to 40 µM in water in a 50 µl volume before LC-MS analysis in negative ionisation mode. Mobile Phase A: 400 mM HFIP and 15 mM TEA in H_2_O; Mobile Phase B: MeOH. Flow rate: 0.5 ml/min. Column temperature: 60 °C. The raw continuum data was deconvoluted to produce zero-charge mass spectra using MassLynx software (Waters).

### Bioinformatics

Publicly available ribosome profiling (riboseq) datasets were analysed using custom in-ioinformatics tools developed to rapidly analyse uORF architecture in any transcript of interest in context of translating ribosomes. Briefly, RefSeq transcript level information were combined with large-scale riboseq datasets including: 46 ribosome footprinting, 8 ribosome initiation, and 34 public mRNA; previously described as part of the GWIPS-viz resource.^15,31^ An aggregation strategy was used to amplify signal and visualize mapped reads in areas with low coverage. Global aggregates of the riboseq datasets were generated per genomic coordinate, and the pyBigWig library was utilized to output a single global bigwig file for each track of interest (Footprints, mRNA, Initiation). SQL queries to the UCSC hg38 database were used to extract sequence features which allow for real-time recalculation and mapping of uORFs and riboseq data on an RNA isoform level.^32^

### Statistics

Statistical analyses were performed using GraphPad Prism v10.1.2 (GraphPad Software, La Jolla, CA, USA).

## Acknowledgements

This work was supported by grants from Great Ormond Street Hospital Sparks Fund/Dravet Syndrome UK and UK MRC (TransNAT) (awarded to MJAW and TCR), the Oxford University Press John Fell Fund and Medical Life Sciences Translational Fund (awarded to TCR). The authors thank Dr Jennifer Frommer for assistance with LC-MS measurements.

