## Supplementary Information for "uORF-targeting steric block antisense oligonucleotides do not reproducibly activate RNASEH1 expression"

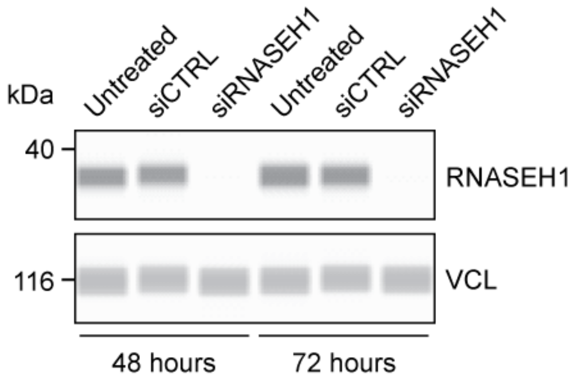


**Figure S1**

**Anti-RNASEH1 antibody validation.**

HeLa cells were transfected with a pool of siRNAs targeting RNASEH1, or a control siRNA pool and protein harvested 48 or 72 hours post transfection. RNASEH1 protein was quantified by Jess capillary western blot. Vinculin (VCL) was used as a loading control. RNASEH1 was detected at the expected size (32 kDa) and was undetectable after siRNA-mediated knockdown.

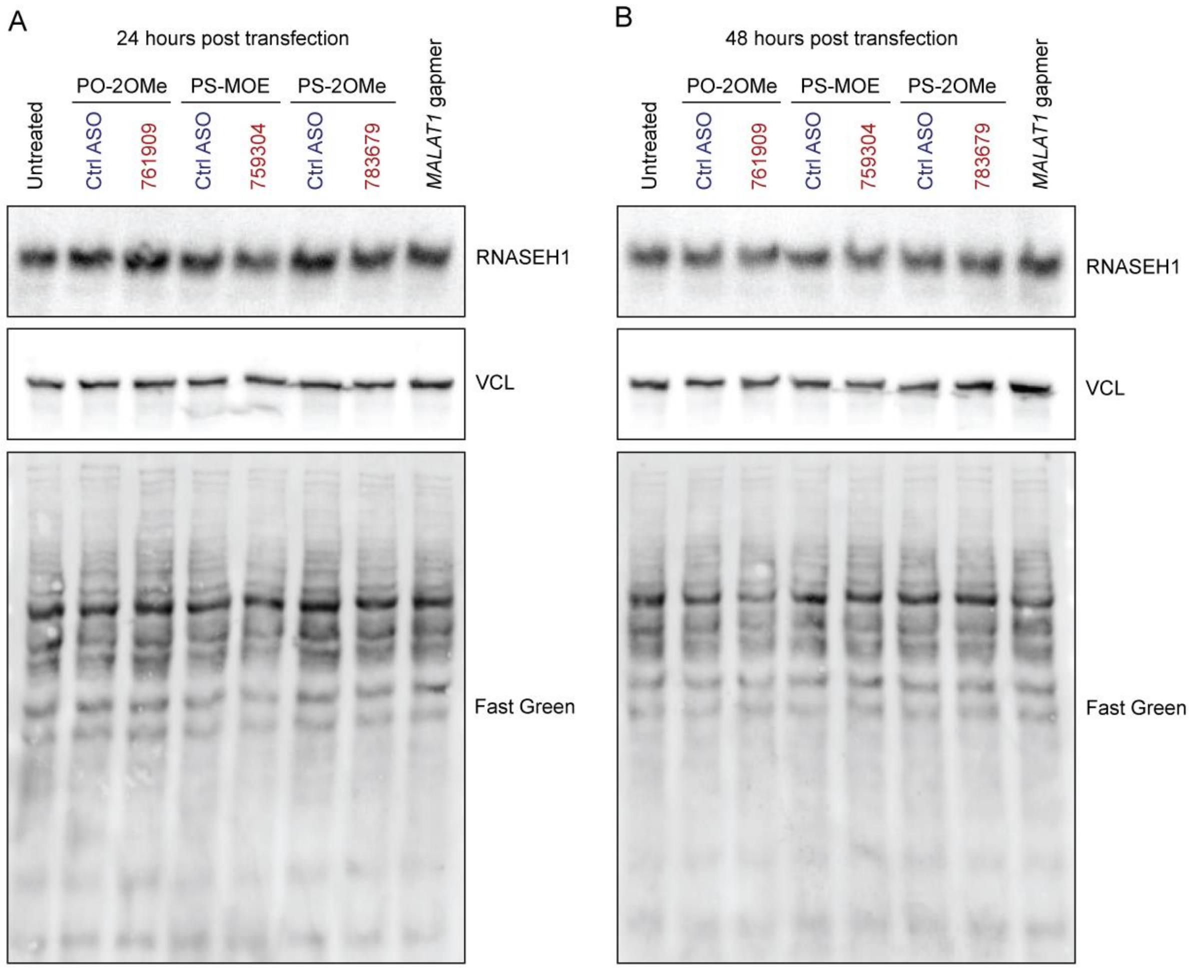


**Figure S2**

**uORF-targeting steric block ASOs do not increase RNASEH1 protein expression at 24 and 48 hours post transfection as assessed by western blot.**

HeLa cells were transfected with ASOs as indicated and protein harvested at (**A**) 24 hours, and (**B**) 48 hours post transfection. Samples were analysed by standard SDS-PAGE western blotting using anti-RNASEH1 antibodies. Vinculin (VCL) was used as a loading control protein, and total protein loading was assessed by Fast Green staining.

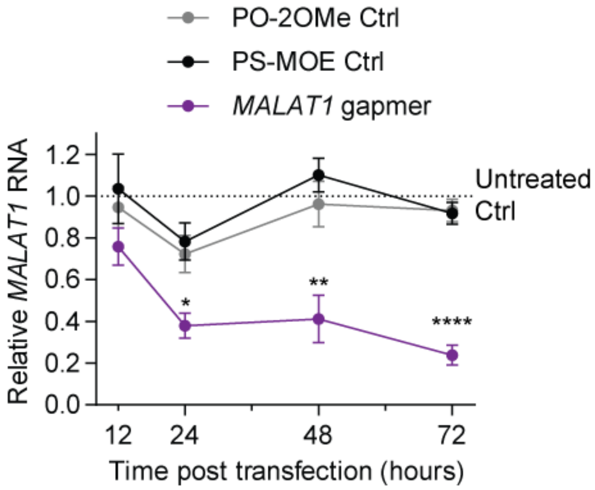


**Figure S3**

**Validation of ASO transfection protocol.**

HeLa cells were transfected with a gapmer (100 nM) ASO targeting *MALAT1* or PO-2OMe and PS-MOE controls and RNA harvested at 12, 24, 48, and 72 hours post transfection. *MALAT1* transcript levels were determined by RT-qPCR and normalised to *RPL10* expression. Values are mean+SEM. Untreated control samples were utilised as calibrator samples and were scaled to a value of 1 at each time point. Statistical differences were determined by one-way ANOVA and Tukey *post hoc* test performed at each time point. *$P$<0.05, **$P$<0.01, ***$P$<0.0001 (comparison of the *MALAT1* gapmer treatment versus the PS-MOE Ctrl), $n$=4 completely independent experiments.

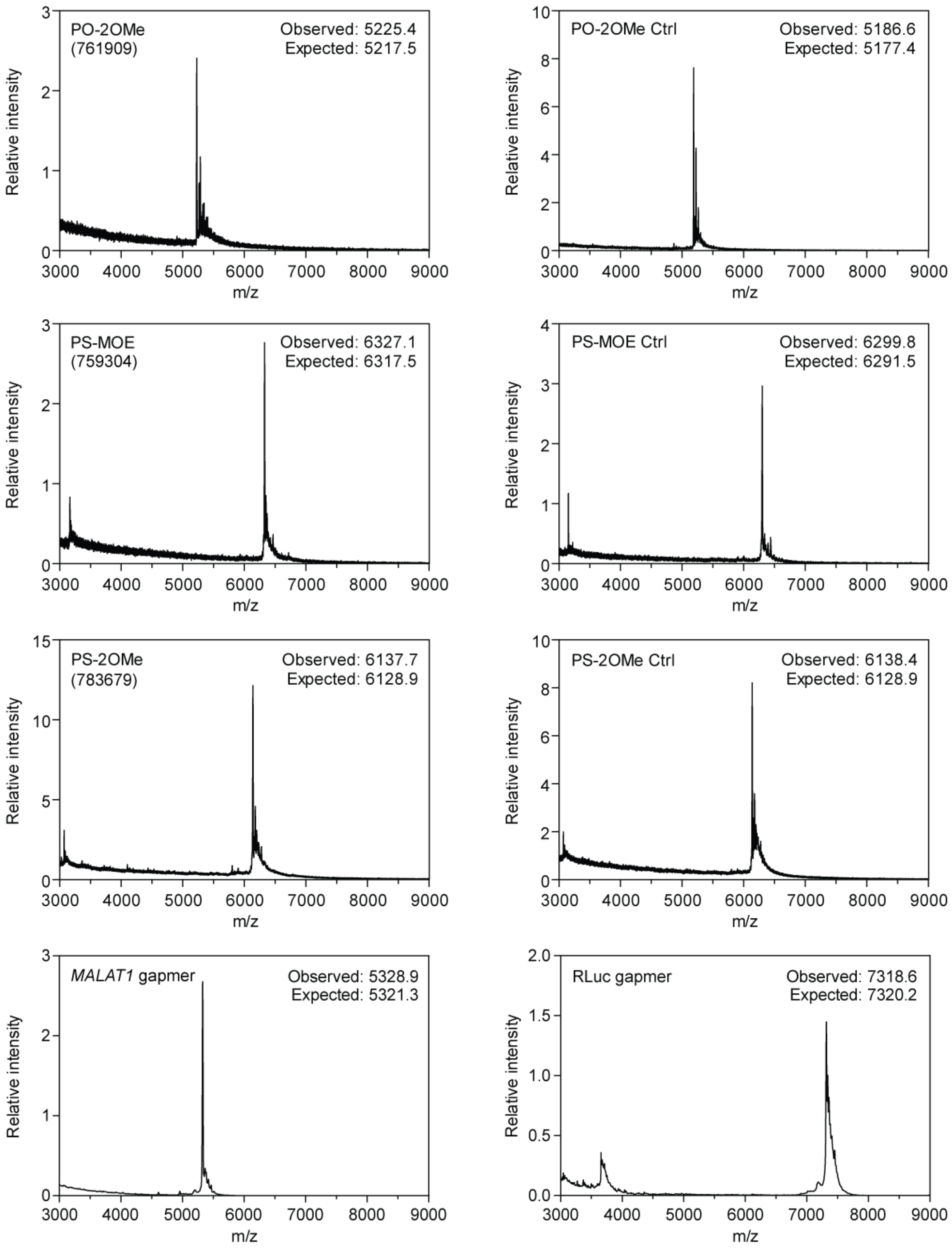
PO-2OMe
(761909)
Observed: 5225.4
Expected: 5217.5
PO-2OMe Ctrl
Observed: 5186.6
Expected: 5177.4
PS-MOE
(759304)
Observed: 6327.1
Expected: 6317.5
PS-MOE Ctrl
Observed: 6299.8
Expected: 6291.5
PS-2OMe
(783679)
Observed: 6137.7
Expected: 6128.9
PS-2OMe Ctrl
Observed: 6138.4
Expected: 6128.9
*MALAT1* gapmer
Observed: 5328.9
Expected: 5321.3
RLuc gapmer
Observed: 7318.6
Expected: 7320.2
Relative intensity
m/z

**Figure S4**

**Confirmation of oligonucleotide integrity by MALDI-TOF-MS.**

MALDI-TOF-MS spectra for oligonucleotides used in this study. Observed and expected mass values are indicated.

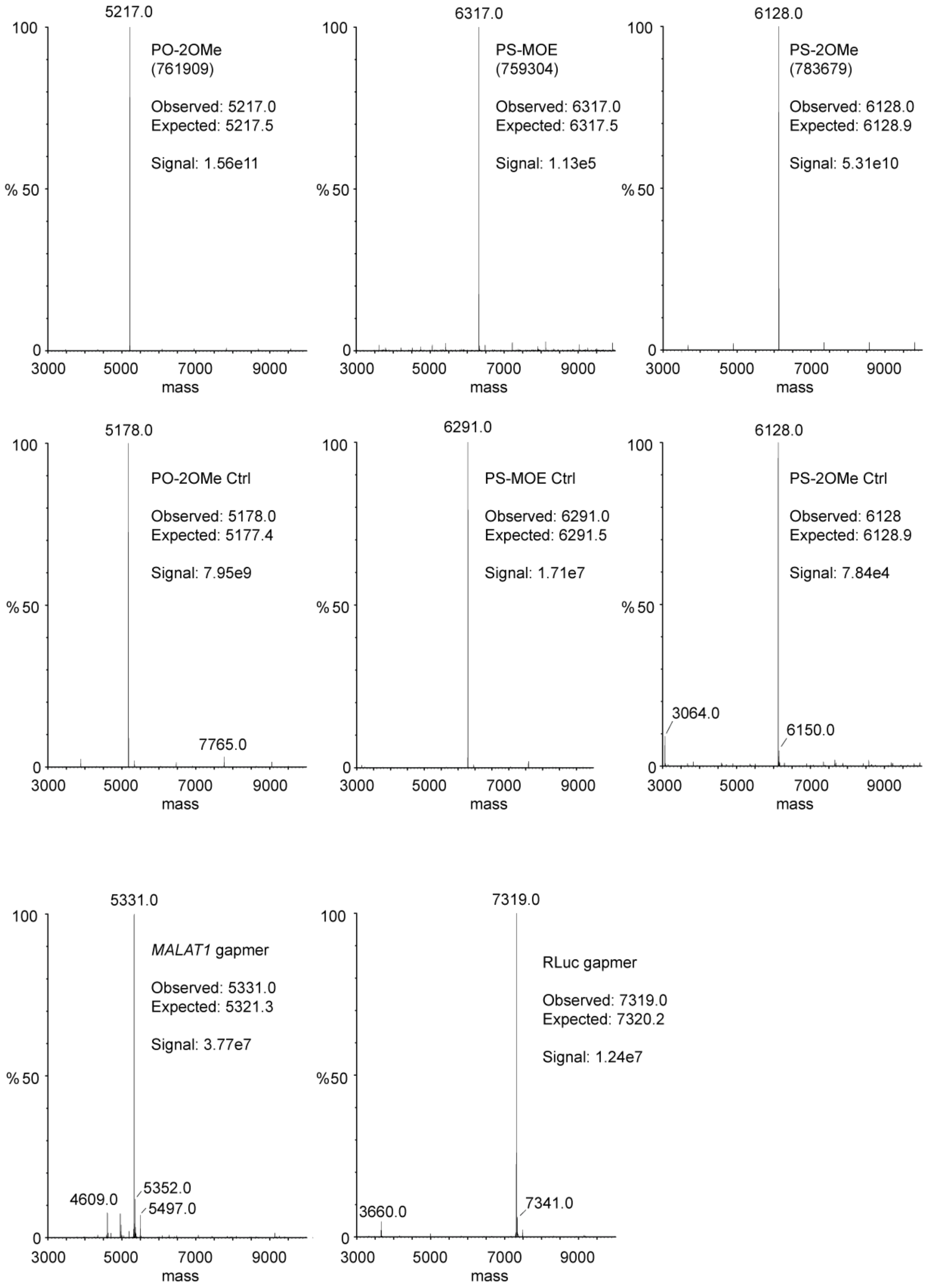
5217.0
PO-2OMe
(761909)
Observed: 5217.0
Expected: 5217.5
Signal: 1.56e11
6317.0
PS-MOE
(759304)
Observed: 6317.0
Expected: 6317.5
Signal: 1.13e5
6128.0
PS-2OMe
(783679)
Observed: 6128.0
Expected: 6128.9
Signal: 5.31e10
5178.0
PO-2OMe Ctrl
Observed: 5178.0
Expected: 5177.4
Signal: 7.95e9
7765.0
6291.0
PS-MOE Ctrl
Observed: 6291.0
Expected: 6291.5
Signal: 1.71e7
6128.0
PS-2OMe Ctrl
Observed: 6128
Expected: 6128.9
Signal: 7.84e4
3064.0
6150.0
5331.0
MALAT1 gapmer
Observed: 5331.0
Expected: 5321.3
Signal: 3.77e7
4609.0
5352.0
5497.0
7319.0
RLuc gapmer
Observed: 7319.0
Expected: 7320.2
Signal: 1.24e7
3660.0
7341.0
100
% 50
0
3000
5000
7000
9000
mass

**Figure S5**

**Confirmation of oligonucleotide integrity by LC-MS.**

Mass spectra for oligonucleotides used in this study as analysed by LC-MS. ASO samples were adjusted to 40 µM in water in a total volume of 50 µl prior to separation. Observed and expected mass values are indicated. Electrospray signal intensity is shown for each spectrum.

| | |
|---|---|
| **RNASEH1-PO2Me (761909)** | |
| mCmAmUmUmUmCmGmAmCmUmCmCmCmGmGmC | CAUUUCGACUCCCGGC |
| **RNASEH1-PO2Me-Control** | |
| mCmGmCmUmUmCmAmCmUmGmCmCmUmAmCmC | CGCUUCACUGCCUACC |
| **RNASEH1-PSMOE (759304)** | |
| /52MOErC/*/i2MOErA/*/i2MOErT/*/i2MOErT/*/i2MOErT/*/i2MOErC/*/i2MOErG/*/i2MOErA/*/i2MOErC/*/i2MOErT/*/i2MOErC/*/i2MOErC/*/i2MOErC/*/i2MOErG/*/i2MOErG/*/32MOErC/ | CATTTCGACTCCCGGC |
| **RNASEH1-PSMOE-Control** | |
| /52MOErC/*/i2MOErG/*/i2MOErC/*/i2MOErT/*/i2MOErT/*/i2MOErC/*/i2MOErA/*/i2MOErC/*/i2MOErT/*/i2MOErG/*/i2MOErC/*/i2MOErC/*/i2MOErT/*/i2MOErA/*/i2MOErC/*/32MOErC/ | CGCTTCACTGCCTACC |
| **RNASEH1-PS2Me (783679)** | |
| mC*mA*mU*mU*mU*mC*mG*mA*mC*mU*mC*mC*mC*mG*mG*mC*mC*mC | CAUUUCGACUCCCGGCCC |
| **RNASEH1-PS2Me-Control** | |
| mC*mG*mC*mU*mU*mC*mA*mC*mU*mG*mC*mC*mU*mA*mC*mC*mG*mC | CGCUUCACUGCCUACCGC |
| ***MALAT1* gapmer** | |
| +C*+T*+A*G*T*T*C*A*C*T*G*A*A*+T*+G*+C | CTAGTTCACTGAATGC |
| **RLuc gapmer** | |
| /52MOErT/*/i2MOErG/*/i2MOErT/*/i2MOErA/*/i2MOErG/*G*A*G*T*A*G*T*G*A*A*/i2MOErA/*/i2MOErG/*/i2MOErG/*/i2MOErC/*/32MOErC/ | TGTAGGAGTAGTGAAAGGCC |

**Table S1**

**Sequences of oligonucleotides used in this study.**

All sequences are written 5′ to 3′. Sequences are provided in IDT notation and as the unmodified sequence only.

| Target | Forward | Reverse |
| --- | --- | --- |
| ***MALAT1*** | GCGTAATGGAAAGTAAAGCCC | CAAACACCTCACAAAACCCC |
| ***RPL10*** | CCTCTTTCCCTTCGGTGTG | AATCTTGGCATCAGGGACAC |

**Table S2**

**RT-qPCR assays used in this study.**

All sequences are written 5′ to 3′.
